# Towards a Unified Framework for Protein Interaction Site Prediction Using Hierarchical Protein Representations

**DOI:** 10.64898/2026.10.18.759950

**Authors:** Viet Thanh Duy Nguyen, John K. Johnstone, Truong-Son Hy

**Affiliations:** Department of Computer Science, the University of Alabama at Birmingham, Alabama, United States

## Abstract

Proteins interact with a wide range of biomolecular partners, yet identifying the interaction sites that mediate these events remains a major challenge for understanding cellular function and guiding drug discovery. Existing computational methods are typically designed for specific interaction classes and often rely on task-specific features. Here, we introduce PRIME-ISP, a structure-based framework for predicting interaction sites across diverse biomolecular interaction types using a unified architecture. PRIME-ISP represents proteins at five organizational levels, including solvent-excluded surfaces, atoms, residues, secondary-structure elements, and whole proteins, and integrates information across these scales through coupled bottom-up and top-down propagation. This hierarchical representation jointly captures local geometry and global structural context, enabling a single architecture to address chemically and geometrically distinct binding problems without task-specific modifications. We evaluate PRIME-ISP on protein, peptide, DNA, RNA, small-molecule, metal, and carbohydrate interaction-site prediction benchmarks. Across these tasks, PRIME-ISP consistently outperforms a recent unified sequence-based predictor and achieves performance comparable to specialized methods while requiring no interaction-specific features. These results demonstrate that hierarchical structural representations provide a flexible foundation for interaction-site prediction across diverse biomolecular interactions. Source code and data are publicly available at https://github.com/HySonLab/PRIME-ISP.

## 1 Introduction

A protein’s biological function is determined largely by where and how it interacts with other molecules. Protein interfaces mediate processes such as signaling and immune recognition, nucleic acid interactions regulate transcription and repair, and binding sites for small molecules, peptides, carbohydrates, and metal ions support catalysis and structural stability. This diversity of interaction modes also creates opportunities to target proteins through different therapeutic modalities. In practice, however, drug discovery has relied predominantly on small molecules, which modulate only a limited fraction of the human proteome and leave many biologically important proteins inaccessible to conventional strategies^1^. This limitation has motivated growing interest in alternative modalities, including proteins, peptides, nucleic acids, and carbohydrates. A protein without a tractable small-molecule pocket may still contain an accessible site for one of these alternative partners^2,3^. Identifying interaction sites across modalities can therefore reveal therapeutic opportunities for targets considered “undruggable” under a single modality. Interaction-site identification is also fundamental to understanding protein function. For example, locating the residues that coordinate a metal ion can help reveal catalytic mechanisms and guide protein engineering^4^. In both functional analysis and therapeutic development, it is not sufficient to determine only whether an interaction occurs. Identifying where it occurs provides the spatial information needed for mutagenesis, molecular docking, and rational design.

Experimental characterization of interaction sites commonly requires techniques such as X-ray crystallography, cryo-electron microscopy, or nuclear magnetic resonance applied to a protein in complex with its binding partner. These experiments remain costly and difficult to perform at proteome scale. Moreover, transient complexes, low-affinity interactions, and proteins that resist structural characterization leave many binding sites experimentally unresolved. Computational methods offer a scalable alternative, but they must account for substantial differences among interaction types. Small molecules commonly bind within compact pockets or cavities, whereas proteins, peptides, and nucleic acids often interact through larger, shape-complementary surface regions. Metal-ion coordination, in contrast, depends on strict geometric and chemical constraints involving a small number of coordinating residues. These differences have led to the development of specialized tools for protein–protein interactions (PPI)^5–8^, protein–peptide interactions (PepPI)^8–11^, protein–DNA interactions (DPI)^12,13^, protein–RNA interactions (RPI)^14–16^, protein–metal interactions (MPI)^4^, protein–carbohydrate interactions (CarbPI)^17^, and protein–small-molecule interactions (SMPI)^18–21^. Most of these methods are designed for only one interaction class. Consequently, analyzing a protein that interacts with several types of partners generally requires multiple specialized and incompatible models.

Despite their biological differences, these prediction tasks share a common computational structure. Given a protein, the objective is to identify the atoms, residues, or surface regions involved in a particular interaction. They can therefore be formulated as a common family of classification or segmentation problems defined over a protein representation. The tasks differ primarily in the interaction class being predicted and the spatial resolution at which an interaction site is represented. This common formulation has motivated recent efforts toward unified prediction. FlexBIP^22^ decouples molecular features from network topology to predict whether two biomolecules interact across a broad range of interaction types. Its objective, however, is to determine whether an interaction exists between two entities rather than to localize the corresponding site within a protein. More closely related to the present problem, ALLSites^23^ predicts interaction sites across a comparably broad range of modalities using a shared sequence-based architecture constructed from pretrained protein language-model embeddings. However, it does not represent three-dimensional protein structure explicitly. This is an important limitation because binding-site geometry is inherently three-dimensional and is expressed through the molecular surface and its underlying atomic organization, neither of which is represented directly by sequence alone.

We take a different route toward unification. Rather than avoiding structural information to achieve generality across interaction modalities, we ask whether protein structure, represented explicitly across multiple physical scales, can provide a common foundation for these tasks. Here, unification refers to the use of a shared architecture and structural representation across interaction classes, with a separate model instance trained for each class. This definition is consistent with ALLSites, which likewise applies the same architecture separately to each modality rather than jointly training a single model across all modalities. This approach is motivated by a simple observation: although these interaction types differ biologically, they are all governed by atom-to-atom physical interactions. A representation constructed directly from protein structure can capture this common physical basis and therefore provides a natural framework for modeling diverse interaction types. To develop such a representation, we build on PRIME^24^, a general hierarchical framework for protein representation learning, and extend it into PRIME-ISP for interaction-site prediction across the seven interaction classes described above. PRIME-ISP represents each protein simultaneously at five levels of physical organization: the molecular surface, atoms, residues, secondary-structure elements, and the protein as a whole. Among these levels, the molecular surface is the region through which most interactions occur, and recent work has shown that surface-based representations are effective for predicting interaction sites and interface properties^25,26^. PRIME-ISP therefore keeps the surface as an explicit level, which lets it model interface geometry directly rather than infer it from sequence. Information is exchanged across all five levels through bottom-up and top-down message passing, rather than processed independently at each scale. This multiscale design allows the same architecture to produce predictions at the resolution each task requires, from atom- or surface-level pocket localization to residue-level classification, without changes to the underlying architecture or a restriction to a single fixed resolution.

To evaluate the generality of PRIME-ISP, we conduct three complementary sets of experiments spanning different interaction classes, spatial resolutions, and binding environments. First, we evaluate residue-level interaction-site prediction across diverse partner classes by comparing PRIME-ISP with a unified sequence-based framework and specialized task-specific methods. This experiment tests whether a shared hierarchical architecture can accommodate interactions with distinct biological and structural characteristics. Second, we evaluate pocket prediction at the atom and molecular-surface levels, examining whether the same representation remains effective at finer spatial resolutions and under a different prediction setting. Third, using proteins with multiple spatially distinct interaction sites, we assess whether interaction-specific models identify partner-dependent binding regions within the same protein structure rather than converging on a generic interaction-prone region. Across these experiments, PRIME-ISP consistently outperforms the unified sequence-based baseline and remains competitive with specialized methods. Its clearest advantages arise for localized binding sites defined by fine-grained geometric and chemical environments. Together, these results show that hierarchical structural representations provide a flexible foundation for predicting interaction sites across diverse partner classes and at residue, atom, and molecular-surface resolutions.

## 2 Results

Across all evaluations in this section, we adopt each benchmark’s original dataset partitions, prediction targets, and evaluation protocol without modification, and compare PRIME-ISP’s performance directly against the baseline results reported in the corresponding source publication rather than reproducing those baselines ourselves. Comparisons therefore assume consistency between our evaluation and the procedure described in each source. Definitions of the metrics used across these benchmarks are given in Section 4.5.

### 2.1 Evaluation on Residue-Level Interaction-Site Prediction Benchmarks

We first evaluate PRIME-ISP on residue-level interaction-site prediction benchmarks spanning seven interaction classes: PPI, PepPI, DPI, RPI, SMPI, MPI, and CarbPI. Except for MPI, we adopt all benchmarks and evaluation settings from ALLSites^23^. The PPI evaluation includes PPI-Train352 with PPI-Test70, PPI-Train9982 with PPI-Test355, and PPI-Train335 with PPI-Test60 and PPI-Test315. The PepPI evaluation includes PepPI-Train1154 with PepPI-Test125 and PepPI-Train640 with PepPI-Test639. The remaining ALLSites benchmarks comprise SMPI-Train1628, SMPI-Valid348, and SMPI-Test348 for SMPI; CarbPI-Train517, CarbPI-Valid129, and CarbPI-Test162 for CarbPI; DPI-Train573 with DPI-Test129 and DPI-Test181 for DPI; and RPI-Train495 with RPI-Test117 for RPI. To extend the evaluation to metal-binding sites, we additionally adopt three ion-specific residue-level benchmarks from MetalBind^4^, corresponding to Ca-, Mn-, and Fe-binding residues, each evaluated independently using the dataset partitions and annotations provided by MetalBind. Together, these benchmarks assess the hierarchical representation’s ability to identify interaction sites ranging from broad protein and nucleic-acid interfaces to localized small-molecule, carbohydrate, and metal-binding environments. ALLSites is a sequence-based method, so its prediction pipeline does not require three-dimensional protein structures. PRIME-ISP, in contrast, requires structural input to construct its hierarchical protein graph. We therefore predict a structure for each protein sequence in the ALLSites benchmarks using ESMFold 2^27^ and use the resulting models to construct the surface-, atom-, residue-, secondary-structure-, and protein-level representations. We retain the original dataset partitions and residue-level annotations without modification.

For each benchmark, we compare PRIME-ISP with ALLSites as the unified sequence-based baseline and, where available,the two strongest task-specific methods reported under the same dataset partition, annotation criteria, and evaluation metric. Candidate methods are drawn from the ALLSites comparisons and more recent literature. The selected task-specific baselines are RGN^28^ for PPI-Test60 and PPI-Test315; EGRET^29^ for PPI-Test70; DELPHI^6^ for PPI-Test355; PepBind^10^ for PepPI-Test639; PepBCL^11^ for PepPI-Test125; EGPDI^30^ for DPI; and GraphBind^15^ for RPI. DeepProSite^8^ is additionally used for PPI-Test60, PPI-Test315, PepPI-Test639, and PepPI-Test125; EnsemPPIS^7^ for PPI-Test70 and PPI-Test355; and DeepPANB^16^ for DPI and RPI. CAPSIF:V and CAPSIF:G^17^ are used for CarbPI. For SMPI, P2Rank^19^ is the only task-specific method with results reported on the same split and is therefore included alongside ALLSites. Because the ALLSites study does not evaluate MPI, we instead compare PRIME-ISP with MetalBind^4^ on the Ca-, Mn-, and Fe-binding benchmarks.

Figure 1 summarizes PRIME-ISP’s residue-level performance across all interaction classes. Compared with ALLSites, the unified sequence-based baseline, PRIME-ISP improves performance across all shared benchmarks, demonstrating the value of explicitly modeling protein structure across multiple physical scales. Its relative performance, however, varies with the nature of the interaction site. For macromolecular interfaces, PRIME-ISP competes with specialized predictors and performs particularly strongly on several PPI benchmarks, though task-specific methods still lead on some protein- and peptide-binding datasets. For nucleic-acid interfaces, PRIME-ISP consistently improves over the sequence-based baseline but remains slightly behind the strongest specialized predictors. The clearest gains occur for localized small-molecule, carbohydrate, and metal-binding sites, where local geometry and chemical organization play a dominant role. We examine these three interaction-site groups in greater detail below.

**Figure 1.**
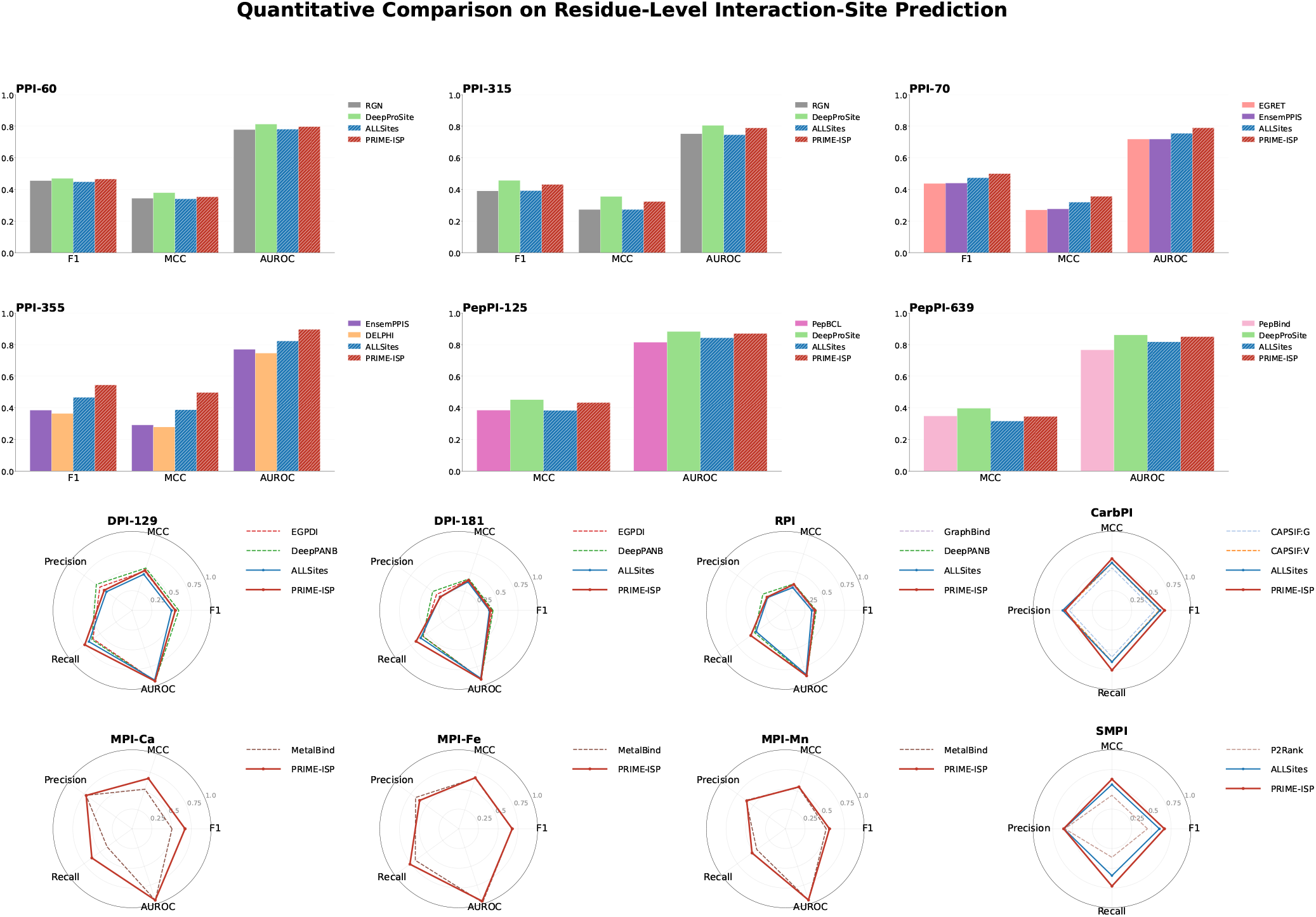
Quantitative comparison between PRIME-ISP and baseline methods on residue-level interaction-site prediction. Results are shown across all seven interaction classes and fourteen benchmark evaluations. Single-class, method-specific baselines are shown with solid bars or thin dashed radar lines, while unified frameworks (ALLSites and PRIME-ISP) are visually distinguished with hatched bars or solid, marker-highlighted radar lines.

#### Macromolecular Interfaces

On PPI-Test60, PPI-Test315, PepPI-Test639, and PepPI-Test125, PRIME-ISP remains competitive with the strongest specialized predictor, although DeepProSite achieves the best reported performance. The difference is small on PPI-Test60, where PRIME-ISP reaches an F1 score of 0.466 compared with 0.470 for DeepProSite, but becomes more pronounced on PPI-Test315 (0.431 versus 0.457). In contrast, PRIME-ISP achieves the highest F1 and MCC among the evaluated methods on PPI-Test70 and PPI-Test355. On PPI-Test355, for example, it reaches an F1 score of 0.545 and an MCC of 0.497, compared with 0.466 and 0.387 for ALLSites. These results show that the hierarchical representation captures substantial interface information using a shared architecture, while the remaining advantage of specialized methods on some benchmarks suggests that broad protein and peptide interfaces may benefit from partner-specific features, conformational information, or interface-specific pretraining.

#### Nucleic-Acid Interfaces

On both DNA- and RNA-binding benchmarks, PRIME-ISP improves over the unified sequence-based baseline but remains below DeepPANB. On DPI-Test129, for example, PRIME-ISP achieves an F1 score of 0.548, compared with 0.501 for ALLSites and 0.592 for DeepPANB. Similarly, its F1 score on RPI improves from 0.336 for ALLSites to 0.374, approaching the 0.391 achieved by DeepPANB. Notably, PRIME-ISP attains higher recall than the competing methods on both tasks, suggesting that the hierarchical representation recovers a larger portion of the binding interface but may also include additional residues outside the annotated region.

#### Localized Binding Sites

The strongest relative performance is observed for small-molecule, carbohydrate, and metal-binding sites. PRIME-ISP increases F1 from 0.601 to 0.664 on SMPI and from 0.609 to 0.665 on CarbPI, outperforming the selected unified and task-specific baselines. It also achieves higher F1 and MCC than MetalBind across all three metal-ion benchmarks, with the largest improvement on Ca-binding sites, where MCC increases from 0.527 to 0.670. These gains are consistent with the multiscale hierarchy: the surface and atomic levels directly capture pocket geometry, accessibility, and local chemical organization, while the coarser levels provide the surrounding structural context.

Overall, the results indicate that hierarchical structural modeling is most effective for compact interaction sites whose identities are strongly constrained by local geometry and chemistry. PRIME-ISP also remains competitive on extended macro-molecular and nucleic-acid interfaces, although these tasks may benefit from explicit partner or conformational information.

The qualitative examples in Figure 2 support this interpretation: predictions form compact regions around localized binding sites and broader coherent regions over extended interfaces, with most errors concentrated near interaction-site boundaries.

**Figure 2.**
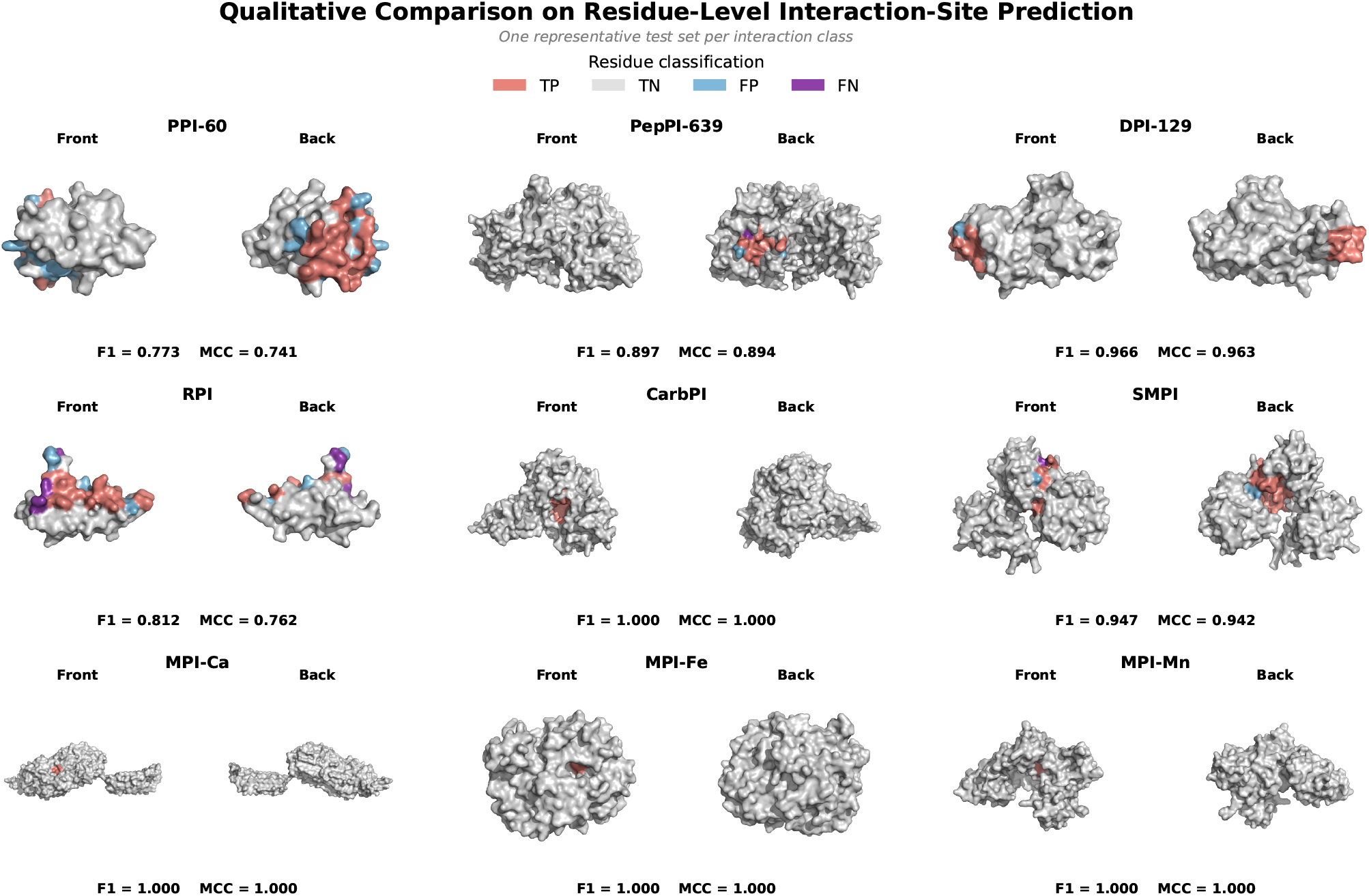
Representative qualitative predictions from PRIME-ISP. For each interaction class, we select one benchmark test set and visualize one representative protein from that set. Predictions are mapped onto the corresponding predicted protein structure and shown from front and back views. Residues are colored by classification outcome (true positive, true negative, false positive, false negative), with the corresponding F1 and MCC scores reported below each pair.

### 2.2 Evaluation on Atom- and Surface-Level Interaction-Site Benchmarks

Residue-level predictions identify which amino acids participate in an interaction but do not resolve which atoms or surface regions actually define the binding pocket, since a labeled residue typically contributes only a subset of its atoms to the interface while the rest face away from it; this distinction matters for applications such as pocket localization, docking, and structure-based molecular design, which require spatial resolution finer than the residue level provides. To evaluate whether PRIME-ISP’s hierarchical representation supports this finer resolution directly, we adopt the protein-ligand binding-site recognition benchmark introduced in SHREC 2022^31^. Unlike the residue-level benchmarks considered in Section 2.1, this benchmark represents proteins at both the atomic and molecular-surface levels, allowing predictions to be evaluated directly on protein atoms and surface vertices. It contains approximately 1,090 protein structures associated with approximately 1,720 experimentally observed ligand-binding sites, partitioned into training and test sets with an approximately 85:15 split. Because a protein may contain multiple co-crystallized ligands, a single structure can have more than one annotated binding site, with up to ten candidate pockets generated and ranked for each test protein. The benchmark provides structures in an anonymized PQR format and evaluates predictions as ranked candidate pockets rather than per-node classifications, both differing from PRIME-ISP’s standard input and output conventions; we adapt PRIME-ISP to this benchmark using the input preprocessing and output pocket-extraction procedures described in Section 4.4.

We compare PRIME-ISP against the baseline methods evaluated on this benchmark by the SHREC 2022 organizers^31^, using the results they report for each participant. At the atomic level, we compare against Fpocket^18^, GNN-Pocket, and NS-Volume^32^,all of which predict pockets directly from atomic coordinates, matching PRIME-ISP’s own atom-level representation. We additionally report DeepSurf^20^, marked as reference-only in Figure 3, since it was pretrained on the external scPDB database and only fine-tuned, rather than trained from scratch, on the SHREC 2022 training set; its results are included for context but are not directly comparable to the other methods. At the surface level, only Point Transformer^33^ among the benchmark’s participants operates on mesh vertices rather than atoms, matching the representation PRIME-ISP uses for its own surface-level predictions, so it serves as the sole comparator reported for that setting.

**Figure 3.**
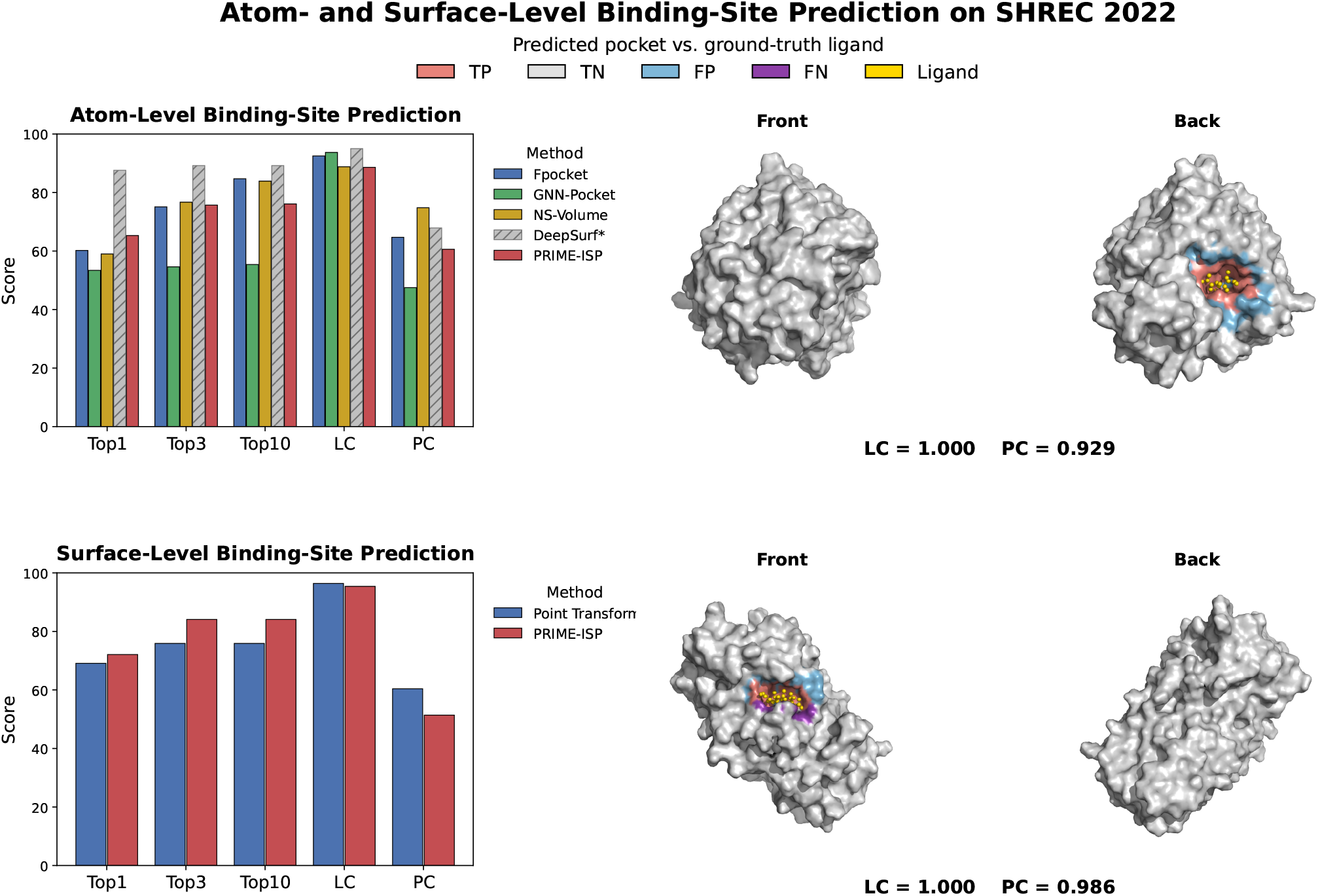
Atom- and surface-level binding-site prediction on SHREC 2022. Top row: atom-resolution methods compared against PRIME-ISP (Atom-level); the hatched DeepSurf* bar denotes a reference point rather than a directly comparable baseline, since it is pretrained on the external scPDB dataset. Bottom row: surface-resolution comparison against Point Transformer. Each row also shows a representative test case, visualizing only the rank-1 predicted pocket against the ground-truth ligand on the protein surface, colored by classification outcome (true positive, true negative, false positive, false negative), with the LC and PC of that pocket reported below each pair.

Figure 3 summarizes the quantitative and qualitative results, reported separately for the atom- and surface-level settings described above. At the atom level, PRIME-ISP reaches Top-1, Top-3, and Top-10 success rates of 65.3%, 75.7%, and 76.1%, outperforming Fpocket, GNN-Pocket, and NS-Volume at Top-1 and remaining competitive with Fpocket and NS-Volume at Top-3, though both surpass it at Top-10. Its LC of 88.6% and PC of 60.6% reflect a trade-off between recovering the ligand-associated region and keeping the predicted pocket tightly localized. DeepSurf reports the strongest atom-level Top-*k* results in this group, consistent with its reference-only status noted above. At the surface level, PRIME-ISP improves Top-1 from 69.1% to 72.1% and both Top-3 and Top-10 from 75.9% to 84.1% relative to Point Transformer, gains of 3.0 and 8.2 percentage points respectively, meaning it more often ranks the correct binding region among its top candidates. Its LC of 95.4% is close to Point Transformer’s 96.4%, but its PC is lower (51.4% vs. 60.4%): PRIME-ISP reliably identifies surface regions in contact with the ligand, but tends to predict a broader pocket that extends beyond it. The representative examples in Figure 3 reflect this pattern: the predicted pocket is concentrated around the ground-truth ligand in both cases, and prediction errors occur mainly near the boundary of the binding region rather than at unrelated locations on the protein surface. Overall, these results show that PRIME-ISP supports direct atom-level pocket prediction and, through the atom-to-surface projection described in Section 4.4, competitive surface-level localization as well.

### 2.3 Case Studies: Partner-Specific Prediction on Multi-Site Proteins

Many pharmacologically important proteins contain multiple, spatially distinct interaction sites for different molecular partners: a small-molecule pocket alongside a separate peptide-, nucleic-acid-, or metal-binding region, for instance. Mapping these partner-specific sites can reveal alternative ways to engage a target when one therapeutic modality is unsuitable. To determine whether PRIME-ISP distinguishes such sites rather than predicting a single generic interaction-prone surface, we examine three proteins with experimentally resolved interaction sites for two distinct partner classes each. To our knowledge, no established benchmark evaluates partner-specific prediction on multi-site proteins under this setting, so we present this analysis as a set of illustrative case studies rather than a systematic evaluation, and the resulting observations should not be interpreted as general performance estimates.

We select thrombin, estrogen receptor *α*, and calmodulin, three pharmacologically important proteins that together illustrate complementary modes of multi-site recognition. Thrombin, a central regulator of blood coagulation and an established anticoagulant target, contains a catalytic pocket targeted by small-molecule inhibitors such as dabigatran, along with spatially distinct exosites recognized by larger partners such as DNA aptamers^34^. Estrogen receptor *α*, an important target in breast cancer and hormone therapy, binds small-molecule agonists and antagonists at its orthosteric pocket while a separate surface, whose accessibility is regulated by ligand binding, is recognized by coactivator proteins. Calmodulin, a central calcium-signaling protein, coordinates Ca^2+^ ions through its EF-hand motifs while binding downstream target peptides through a distinct hydrophobic groove^35^. Together, these three targets let us test whether interaction-specific predictors identify different regions of the same protein depending on the selected partner class.

For each target, we select a single experimentally resolved complex containing two partner classes bound to the same protein structure. This avoids potential inconsistencies in construct boundaries, residue numbering, and protein conformation that could arise when combining annotations from separate structural depositions. The selected complexes are thrombin bound to a DNA aptamer and a small-molecule inhibitor (PDB 8TQS^34^), estrogen receptor *α* bound to a small-molecule agonist and a coactivator peptide (PDB 3L03), and calmodulin bound to four Ca^2+^ ions and a target peptide (PDB 1CDL^35^). Together, these complexes cover four interaction classes: SMPI, PepPI, DPI, and MPI. For each complex, all bound partners, water molecules, and crystallization additives are removed, while the target protein coordinates and residue numbering are retained. The resulting partner-free structure is then provided independently to the corresponding interaction-class-specific models. Specifically, thrombin is evaluated using the SMPI and DPI models, estrogen receptor *α* using the SMPI and PepPI-Train1154 models, and calmodulin using the MPI and PepPI-Train1154 models. We directly reuse the trained model checkpoints employed in the residue-level benchmark evaluation in Section 2.1, without additional training, fine-tuning, or hyperparameter selection on the case-study proteins. Although these models share the same hierarchical architecture and input-feature construction, each is trained separately on its corresponding interaction-class benchmark and therefore has distinct learned parameters. To ensure consistency with the labels used to train each interaction-class-specific model, ground-truth sites for the case studies are constructed using the same distance cutoffs as the corresponding training datasets. Specifically, a residue is labeled as interacting if any of its atoms lies within 6.5 Å of a small-molecule ligand atom, 5.0 Å of a peptide or nucleic-acid partner atom, or 3.0 Å of a bound metal ion^4,23^. Ground-truth labels are constructed independently for each partner in the original complex, and each model prediction is evaluated only against the labels for its corresponding interaction class. This design assesses whether independently trained PRIME-ISP models produce distinct, partner-class-specific prediction maps from the same protein structure rather than converging on a generic interaction-prone region.

As shown in Figure 4, PRIME-ISP predicts five of the six sites with high precision, with the strongest overall performance on the calmodulin metal-ion site. The calmodulin/MPI site is recovered with near-perfect accuracy (precision = 0.952, recall = 1.000, F1 = 0.976), correctly identifying all four calcium-coordinating loops with almost no error. The small-molecule sites are similarly reliable: thrombin/SMPI (precision = 0.870) and ER*α*/SMPI (precision = 0.750) both localize the correct pocket with few false positives, as do the peptide sites, ER*α*/PepPI (F1 = 0.741) and calmodulin/PepPI (precision = 1.000). Across all six sites, precision is consistently high (five of six at precision ≥0.75), meaning PRIME-ISP rarely flags a residue outside the true site; where it falls short is completeness, often catching the core of a binding region without its full extent, as with calmodulin/PepPI (recall = 0.204), where the true site forms one contiguous groove and the model correctly identifies its center but leaves the flanking residues below threshold. The one case where both precision and recall are low is thrombin/DPI (precision = 0.250, recall = 0.094), where PRIME-ISP fails to localize the DNA aptamer site. A likely explanation lies in the training distribution: the DPI benchmark consists predominantly of natural DNA-binding proteins, such as transcription factors and DNA-processing enzymes, whose interfaces evolved to recognize DNA. Thrombin is not a natural DNA-binding protein; the aptamer instead binds an exosite that evolved to recognize protein partners, so the structural and physicochemical features of this site may differ substantially from the DNA-binding interfaces the predictor learned. This points to a limitation of the current approach: predictions are conditioned on the partner’s interaction class rather than on the identity of the specific partner, so a protein engaged by an atypical partner within a class, such as an engineered aptamer, can be predicted less reliably than a protein that binds its natural partner.

**Figure 4.**
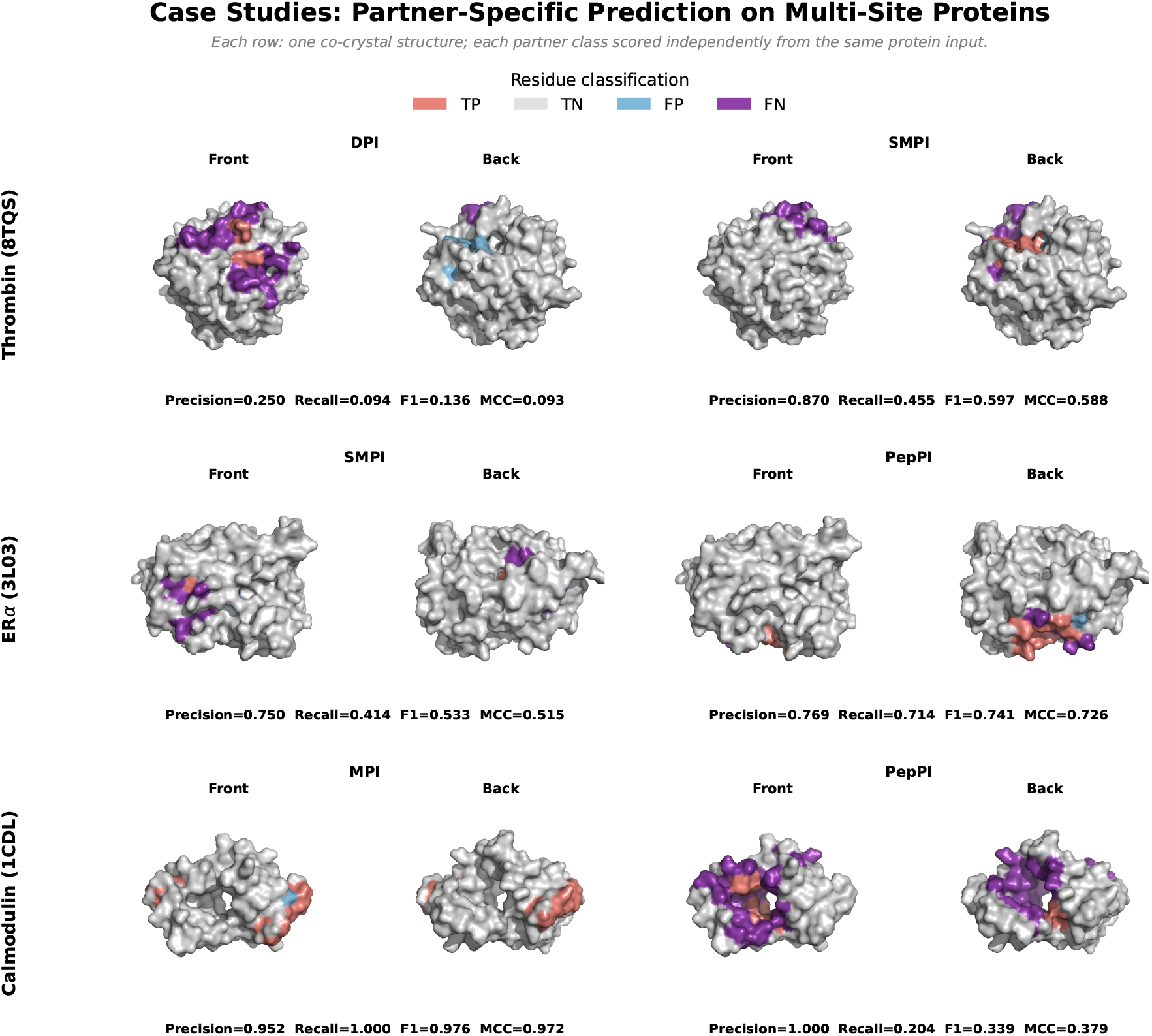
Partner-specific PRIME-ISP predictions on three case studies. Each row is one experimental complex (thrombin, ER*α*, calmodulin); each pair of columns is one partner class scored independently on that same partner-free structure, shown from the front and back. The surface is colored by residue classification (true positive, true negative, false positive, false negative). Precision, recall, F1, and MCC below each pair are computed for that specific site against the experimentally observed binding residues.

### 2.4 Ablation Studies

To evaluate the principal architectural components of PRIME-ISP, we conduct two complementary ablation studies on the SMPI benchmark. First, we examine the contribution of individual structural levels by removing selected levels from the hierarchical graph. Second, we evaluate the directionality of cross-level communication by comparing the complete bidirectional architecture with bottom-up-only and top-down-only variants. We select SMPI because small-molecule binding pockets are influenced by local surface geometry and atomic chemistry, while residue-, secondary-structure-, and protein-level representations provide progressively broader structural context. All variants are trained and evaluated using the same dataset partitions, input features, optimization settings, and evaluation procedures, with performance measured on SMPI-Test348.

#### 2.4.1 Contribution of Hierarchical Representation Levels

Starting from the complete hierarchy, we remove the surface, atomic, secondary-structure, and protein levels in separate experiments while retaining the residue level from which predictions are generated. Because the assignment operators **Π**^(*l*)^ connect only adjacent levels, removing an intermediate level also disconnects any levels that depend on it for communication with the residue level. Specifically, removing the atomic level disconnects the surface level, whereas removing the secondary-structure level disconnects the protein level. In contrast, removing the surface or protein level isolates the contribution of a single level because these levels occupy the two ends of the hierarchy.

Table 1 reports the results. Removing the atomic and surface levels produces the largest degradation, reducing F1 from 0.664 to 0.642 and MCC from 0.628 to 0.604, corresponding to relative decreases of 3.3% and 3.8%, respectively. Because this variant removes both levels, the observed degradation reflects the combined loss of atomic chemistry and surface geometry. Removing the surface level alone, by contrast, results in substantially smaller decreases of 0.8% in F1 and 1.0% in MCC, suggesting that atomic information accounts for most of the benefit provided by the two finest-grained levels on SMPI. Removing the secondary-structure and protein levels produces relative decreases of 1.4% in F1 and 1.4% in MCC, indicating that structural context coarser than the residue level also contributes to pocket recognition, while removing only the protein level has a smaller effect, reducing F1 by 0.5% and MCC by 0.6%. Together, these results suggest that secondary-structure organization provides useful intermediate-scale context, whereas the global sequence representation contributes a smaller, complementary signal. Although some ablated variants achieve slightly higher recall or AUROC, the complete hierarchy obtains the highest F1, MCC, and precision, reflecting the most balanced overall performance.

**Table 1.** Ablation of hierarchical representation levels on SMPI-Test348. Each row removes the indicated level or levels while retaining the residue level used for prediction. Removing the atomic level also disconnects the surface level, whereas removing the secondary-structure level also disconnects the protein level. The best value in each column is shown in bold.

| Configuration | F1 | MCC | Precision | Recall | AUROC |
| --- | --- | --- | --- | --- | --- |
| Full hierarchy | **0.664** | **0.628** | **0.612** | 0.726 | **0.934** |
| w/o surface | 0.659 | 0.622 | 0.595 | 0.737 | **0.934** |
| w/o atom (and surface) | 0.642 | 0.604 | 0.577 | 0.725 | 0.932 |
| w/o secondary structure (and protein) | 0.655 | 0.619 | 0.582 | **0.749** | 0.933 |
| w/o protein | 0.661 | 0.624 | 0.601 | 0.734 | 0.932 |

#### 2.4.2 Contribution of Bidirectional Cross-Level Communication

Having established that the intermediate levels contribute complementary information, we next ask whether the *direction* of communication between them matters equally. The complete model alternates bottom-up aggregation, which transfers fine-grained geometric and chemical information toward coarser levels, with top-down refinement, which broadcasts global and intermediate-scale context toward finer levels. We compare this bidirectional architecture with a bottom-up-only variant in which top-down refinement is disabled and a top-down-only variant in which bottom-up aggregation is disabled. All hierarchical levels, within-level message-passing operations, node features, and prediction heads are retained across the three configurations, ensuring that the variants differ only in the direction of cross-level information propagation.

We summarize these results in Table 2. The bidirectional configuration achieves the best F1, MCC, precision, and AUROC, confirming that bottom-up and top-down propagation carry complementary information rather than one direction subsuming the other. Removing either direction produces a similar drop in overall performance, with MCC falling by 0.019 without top-down refinement and by 0.023 without bottom-up aggregation, suggesting that the two directions contribute comparable amounts of information on this task rather than one being clearly more important. The single-direction variants also show a systematic precision-recall trade-off relative to the full model: bottom-up-only reaches the highest recall (0.736) but the lowest precision (0.577), while top-down-only sits closer to the bidirectional model on precision but recovers fewer true binding residues. This pattern is consistent with the role each direction plays in the architecture. Without top-down refinement, coarser-level context never propagates back down to correct residue-level predictions, so the model tends to over-predict, flagging more residues as positive at the cost of specificity. Without bottom-up aggregation, residue-level predictions are made without the fine-grained geometric and chemical signal that bottom-up propagation would have supplied, so the model becomes more conservative but misses more true positives. The bidirectional model balances these effects, retaining nearly all of the bottom-up variant’s recall (0.726 vs. 0.736) while achieving higher precision than either single-direction variant (0.612 vs. 0.577 and 0.590), which yields its higher F1 and MCC.

**Table 2.** Ablation of cross-level message-passing directionality on SMPI-Test348. All configurations use the same hierarchical levels, node features, and within-level message-passing modules. The best value in each column is shown in bold.

| Configuration | F1 | MCC | Precision | Recall | AUROC |
| --- | --- | --- | --- | --- | --- |
| Bottom-up only | 0.647 | 0.609 | 0.577 | **0.736** | 0.929 |
| Top-down only | 0.644 | 0.605 | 0.590 | 0.708 | 0.928 |
| Bidirectional | **0.664** | **0.628** | **0.612** | 0.726 | **0.934** |

## 3 Discussion

Accurate interaction-site prediction requires representations that capture both the local environments governing molecular recognition and the broader structural context in which they occur. We developed PRIME-ISP to address this need by organizing the molecular surface, atoms, residues, secondary-structure elements, and the complete protein into a physically grounded hierarchy. Deterministic relationships and bidirectional communication between adjacent levels allow the framework to integrate fine-grained geometry and chemistry with progressively broader structural context, providing a common architecture for interaction-site prediction across partner classes and at residue, atom, and surface resolutions.

The benchmark results demonstrate that hierarchical structural modeling provides information complementary to sequence-based representations. PRIME-ISP consistently outperforms the unified sequence-based baseline and remains competitive with specialized methods. Its clearest advantages occur for localized small-molecule, carbohydrate, and metal-binding sites, whose recognition depends strongly on pocket geometry and local atomic organization. The smaller gains on protein, peptide, DNA, and RNA interfaces suggest that broader interfaces depend more strongly on distributed surface properties, conformational flexibility, and partner-specific context. The SHREC 2022 results further show that the same hierarchy supports direct atom-level prediction and surface-level pocket localization, demonstrating its applicability beyond residue classification.

The multi-site case studies provide an exploratory view of how separately trained interaction-specific models behave on the same protein structure. The models produce distinct maps for different partner classes and recover several experimentally observed sites, including the Ca^2+^-coordination sites of calmodulin and the small-molecule and peptide-binding regions of estrogen receptor *α*. Several predictions exhibit high precision but lower recall, indicating that PRIME-ISP often identifies a reliable core of an interaction site without recovering its complete extent. The thrombin DNA-aptamer interface is the most challenging case. Because PRIME-ISP predicts general interaction propensity for a partner class without receiving the identity or structure of the specific partner, an atypical synthetic aptamer may bind differently from the natural protein-DNA interfaces represented in the training data. This result highlights the distinction between predicting a general DNA-binding region and predicting the site used by a particular DNA molecule.

The ablation studies support the importance of both the hierarchical levels and their communication. Removing the atomic and surface levels causes the largest performance reduction on small-molecule interaction-site prediction, confirming the contribution of fine-grained chemical and geometric information beyond residue-level features. The secondary-structure and protein levels provide smaller but complementary gains through broader structural context. Neither bottom-up nor top-down communication alone matches the full model: bottom-up propagation supplies fine-grained surface and atomic information, whereas top-down refinement provides secondary-structure and protein-level context. Their combination achieves the strongest overall performance, demonstrating that the two directions contribute complementary information.

A primary direction for future work is to condition interaction-site prediction explicitly on the interacting partner. The current formulation uses a separately trained model for each interaction class and estimates where a protein is generally likely to interact with that class, but it does not directly account for the properties of a specific partner. Future models could combine the hierarchical protein representation with molecular graphs for small molecules, sequence or structural representations for proteins and peptides, nucleic-acid representations for DNA and RNA, and element- or coordination-specific features for metal ions. Cross-attention or geometric interaction modules could then identify the protein region most likely to bind a specified partner. Joint multitask training could further replace the separate interaction-specific models with a shared encoder and partner-aware prediction heads, allowing common structural principles to be learned across interaction classes. Finally, because no established benchmark currently evaluates multiple partner-class-specific sites on a common protein structure, a systematically curated multi-site benchmark would enable a more comprehensive assessment of partner-conditioned prediction. Together, these developments could extend PRIME-ISP from predicting general interaction propensity for a partner class toward identifying the binding site of a specific interacting molecule.

## 4 Methods

### 4.1 Problem Formulation

Let a protein structure be represented as a hierarchical graph

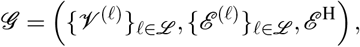

where ℒ = {surface, atom, residue, secondary structure, protein} denotes the set of structural levels corresponding to solvent-excluded surface points, atoms, residues, secondary-structure elements, and the complete protein, respectively. For each level *l* ∈ ℒ, *V* ^(*l*)^ is the associated node set, and *ℰ* ^(*l*)^ is the set of within-level edges. The set *ℰ* ^H^ contains cross-level edges that encode hierarchical relationships between nodes at adjacent structural levels.

Each node 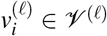 is associated with an input feature vector 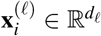, which describes the sequence, geometric, structural, or physicochemical properties available at level *l*. Within-level edges encode relationships appropriate to the corresponding structural scale, such as spatial proximity, covalent connectivity, sequence adjacency, or secondary-structure organization. Cross-level edges associate fine-grained nodes with their corresponding coarse-grained structural units, such as atoms with residues, residues with secondary-structure elements, and secondary-structure elements with the complete protein.

Let *j* = {protein, peptide, DNA, RNA, small molecule, metal, carbohydrate} denote the set of interaction partner classes considered in this work. For a task associated with partner class *τ* ∈*j*, let *l*_*τ*_ ∈ ℒ denote the structural level at which predictions are evaluated. The corresponding set of prediction units is 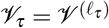.

For each prediction unit 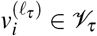, we define a binary interaction-site label

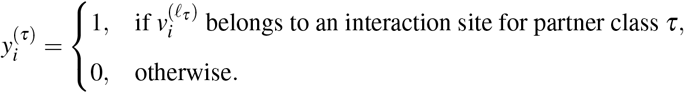

A node is considered part of an interaction site according to the annotation rule defined by the corresponding benchmark. Depending on the task, this rule may be based on the minimum distance to the interacting partner, overlap with an annotated binding region, or another benchmark-specific geometric criterion.

For each interaction partner class *τ* ∈*j*, we train a separate instance of the same hierarchical architecture, with its own learned parameters ***θ*** _*τ*_, to assign an interaction probability to each task-level prediction unit given the protein graph *G* :

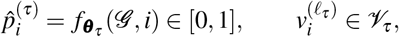

where 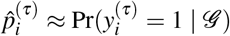 is the estimated probability, under the class-*τ* model, that prediction unit 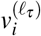 belongs to an interaction site of type *τ*. All |*j*| models share the same hierarchical architecture, node feature definitions, and training procedure, but each ***θ*** _*τ*_ is fit independently to its own benchmark; no parameters are shared across partner classes at inference time.

A binary prediction is subsequently obtained using a decision threshold *δ*_*τ*_ :

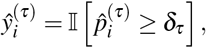

where I[·] denotes the indicator function. Following common practice, we fix *δ*_*τ*_ = 0.5 for all tasks.

Under this formulation, PPI, PepPI, DPI, RPI, SMPI, MPI, and CarbPI site prediction are treated as instances of a common hierarchical node-classification problem, each solved by its own class-specific model 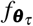. These tasks differ in four principal aspects: the structural level at which predictions are evaluated, the definition of a positive interaction-site label, the size and distribution of the available data, and the benchmark-specific evaluation protocol. The hierarchical representation, node feature definitions, and core model architecture remain unchanged across tasks, except for the prediction head and output resolution required by each benchmark; what differs across tasks is the learned parameters ***θ*** _*τ*_ themselves, each optimized separately on its own benchmark.

### 4.2 PRIME-ISP: A Hierarchical Framework for Interaction-Site Prediction

We introduce PRIME-ISP, a structure-based framework for predicting protein interaction sites across diverse partner classes and structural resolutions. As illustrated in Figure 5, PRIME-ISP builds on PRIME^24^, a multiscale protein representation learning framework that represents a protein as a hierarchy of five physically grounded structural levels: the molecular surface, atoms, residues, secondary-structure elements, and the complete protein. Adjacent levels are connected through deterministic assignments derived from the protein’s geometric and biochemical organization, rather than treated as independent modalities. Within this hierarchy, bottom-up aggregation propagates fine-grained geometric and chemical information toward coarser levels, while top-down refinement communicates global structural context back to finer levels. PRIME-ISP adapts this architecture to interaction-site prediction by applying a node-level prediction head to the structural level required by each task. The prediction head assigns an interaction probability to each node at the selected level, producing a localized interaction-site prediction while preserving the same hierarchical encoder and input representations across interaction classes.

**Figure 5.**
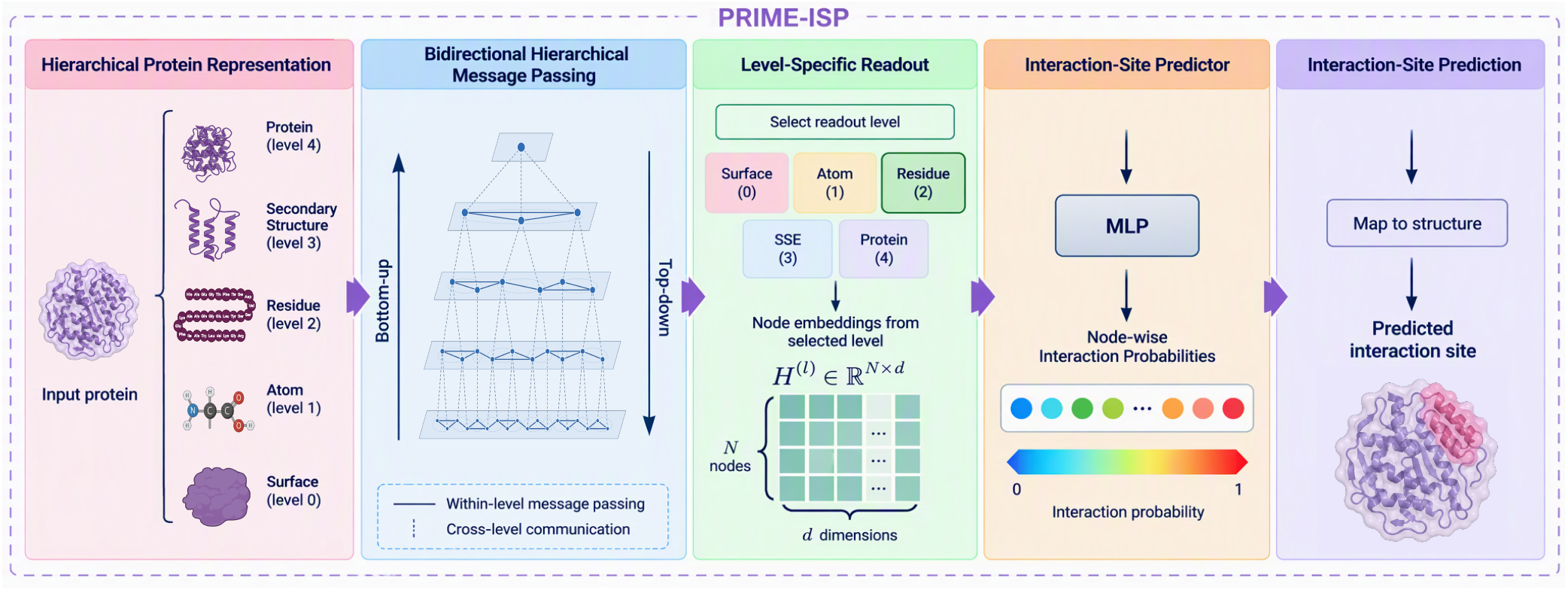
Overview of the PRIME-ISP framework. An input protein is represented as a five-level hierarchy comprising the molecular surface, atoms, residues, secondary-structure elements, and the complete protein. Within-level message passing updates nodes at each structural resolution, while bottom-up aggregation and top-down refinement exchange information between adjacent levels. The node embeddings at a selected readout level form a matrix **H**^(*l*)^ ∈ ℝ^*N*×*d*^, where *N* is the number of nodes at that level, and *d* is the embedding dimension. A prediction head maps each node embedding to an interaction probability, producing a node-wise probability map over the selected structural level. These probabilities are mapped onto the protein structure to localize the predicted interaction site.

#### Hierarchical graph construction

Given an input protein structure, PRIME-ISP builds a hierarchy *G* ordered from fine to coarse resolution. Atomic and residue nodes are taken directly from the protein’s PDB structure. At the surface level, the solvent-excluded molecular surface is computed with PyMOL^36^ and represented as a triangular mesh, with each face treated as a node positioned at its centroid. At the atomic level, nodes correspond to protein heavy atoms; at the residue level, to amino acid residues; and at the secondary-structure level, to contiguous helices, strands, and loops identified from the backbone using DSSP algorithm^37^. The coarsest level contains a single node representing the complete protein.

Edges at the finest, surface level are constructed via *k*-nearest neighbors (*k* = 8) over face centroids, connecting each face to its spatially closest neighboring faces; this yields an adjacency matrix *A*^(0)^. Connectivity at each coarser level *l* is then derived deterministically from *A*^(0)^ by successive application of the assignment matrices,

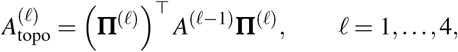

so that two coarse-level nodes are connected whenever their constituent fine-scale nodes are connected at the level below; we refer to 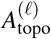 as the *hierarchy-derived topological edges* at level *l*, since they are inherited from surface connectivity rather than computed directly from atomic or residue coordinates.

Because the surface mesh covers only the solvent-exposed exterior of the protein, structural units with no assigned surface faces, such as atoms and residues buried in the protein core, would remain disconnected from the rest of the graph under 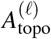 alone. To ensure that every atom and residue participates in message passing regardless of burial depth, we complement the topological edges at the atomic and residue levels with spatial *k*-nearest-neighbor edges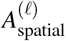, computed directly from three-dimensional coordinates (*k* = 8) independently of surface-derived connectivity. Rather than merging the two edge sets into a single adjacency, we retain them as two distinct relations and process them with a relational graph convolution: neighbor aggregation is computed separately for topological and spatial neighbors, each with its own learnable linear transformation, and the two resulting messages are summed before the residual update. This keeps hierarchy-derived connectivity and direct spatial proximity as separable signals rather than conflating them into a single, ambiguous notion of adjacency. Both relation-specific adjacency matrices are normalized independently using symmetric degree normalization, 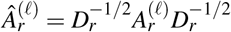 for *r*∈{topo, spatial}, which prevents high-degree nodes from dominating aggregation and stabilizes training. At the secondary-structure and protein levels, where every structural unit derives its identity directly from the residue-level partition rather than from surface-derived coarsening, only the single topological adjacency is used.

Adjacent levels are linked through deterministic assignment operators: each surface face is assigned to its nearest heavy atom by Euclidean distance between the face centroid and atomic coordinates, each atom to its parent residue via covalent bonding structure, each residue to its secondary-structure element, and every secondary-structure element to the protein-level node. Let 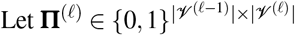 denote the assignment matrix from level *l* − 1 to level *l*:

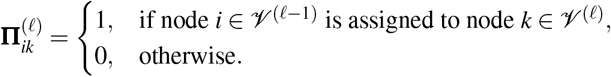

These matrices encode fixed, known geometric and biochemical relationships and are not updated during training. We note that the surface-to-atom assignment does not guarantee every atom receives a face; such atoms remain connected within their level via the spatial edges above, and still participate in cross-scale fusion through the atom- and residue-level partition matrices, which are defined independently via covalent bonding and DSSP segmentation rather than surface proximity.

#### Node representations

To isolate the contribution of the hierarchical architecture from that of pretrained components, PRIME-ISP omits the pretrained surface and atomic encoders used in the original PRIME framework. The surface and atomic levels are initialized entirely from handcrafted geometric and physicochemical features; the residue level combines handcrafted descriptors with contextual ESM-2^38^ embeddings; the secondary-structure level uses handcrafted segment descriptors; and the protein-level node is initialized with a global ESM-2 representation of the full sequence. Input features at each level are defined as follows.

- **Surface level**. Each triangular face is represented by a seven-dimensional handcrafted feature vector consisting of face area, distance from the protein centroid, discrete mean curvature, discrete Gaussian curvature, local concavity, average edge length, and normalized aspect ratio. Concavity is measured from the alignment between the face normal and the outward radial direction from the protein centroid. The normalized aspect ratio is

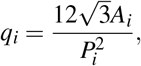

where *A*_*i*_ and *P*_*i*_ are the area and perimeter of face *i*; *q*_*i*_ = 1 for an equilateral triangle and decreases as the face becomes less regular. As with all surface features, *q*_*i*_ is standardized across the faces of each protein.
- **Atomic level**. Each heavy atom is represented by a 17-dimensional vector combining chemical, residue-level, and local geometric information: a six-dimensional element-type encoding, a backbone indicator, and van der Waals radius; the hydrophobicity and charge of the parent residue; binary indicators for hydrogen-bond donor, hydrogen-bond acceptor, and aromatic-residue membership; and four local geometric descriptors, normalized distance to the protein centroid, nearest-neighbor distance, and buriedness within 5 Å and 8 Å.
- **Residue level**. Each residue is represented by a 1,318-dimensional vector. Handcrafted components include a 20-dimensional amino acid one-hot encoding; three backbone geometric descriptors (bond angle, improper torsion, dihedral angle); ten physicochemical descriptors (solvent accessibility, hydrophobicity, hydrophilicity, molecular volume, isoelectric point, helix/sheet/coil propensities, residue charge, and atom count); and backbone *φ* /*ψ* torsion angles as sine-cosine pairs together with relative solvent accessibility. Each residue is additionally assigned a 1,280-dimensional contextual embedding from layer 33 of ESM-2 650M, with beginning- and end-of-sequence tokens excluded.
- **Secondary-structure level**. Each element is represented by a four-dimensional vector: a three-way one-hot encoding of structural class (loop, helix, strand) and its normalized length in residues.
- **Protein level**. The complete protein is represented by the 1,280-dimensional beginning-of-sequence (BOS) embedding from layer 33 of ESM-2 650M, used instead of mean-pooled residue embeddings to avoid duplicating residue-level information already aggregated through bottom-up message passing.

All continuous handcrafted features are standardized using training-set statistics. Because feature dimensionality differs across levels, each input matrix is projected into a common hidden dimension by a level-specific learnable transformation:

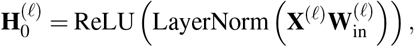

where 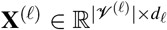 is the input feature matrix at level *l*, 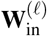 is a level-specific learnable projection, and 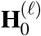 is the resulting initial hidden representation.

#### Multiscale message passing

PRIME-ISP learns context-aware node representations through repeated hierarchical message-passing layers, each combining within-level message passing, bottom-up aggregation, and top-down refinement.

Within-level message passing first refines representations at each level independently through a graph convolution block with pre-layer normalization, followed by a two-layer feedforward network with GELU activation. At the atomic and residue levels, where two relations are defined, neighbor aggregation is computed separately over the topological and spatial adjacency matrices introduced above, each using its own relation-specific weight matrix, following a relational graph convolution,

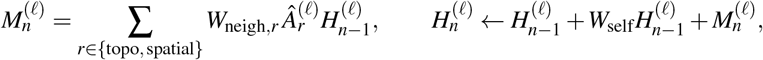

so that the model distinguishes hierarchy-derived connectivity from direct three-dimensional proximity rather than treating them as a single, undifferentiated notion of adjacency. At the remaining levels, where only the single topological adjacency is defined, neighbor aggregation reduces to the *r* = topo term alone.

Bottom-up propagation then transfers fine-grained information to the adjacent coarser level, from level *l* − 1 to level *l*:

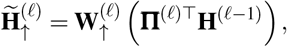

with learnable projection 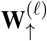. Applied successively, this propagates surface geometry to atoms, atomic information to residues, residue information to secondary-structure elements, and secondary-structure information to the protein level.

Top-down refinement broadcasts coarse-level context back to the adjacent finer level:

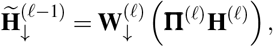

with learnable projection 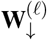. This lets global and intermediate-scale context refine finer-resolution predictions: a residue’s representation can incorporate both local atomic and surface detail from below, and secondary-structure and protein-level context from above.

Context received from an adjacent level is integrated by gated fusion. Given a current representation **z** and cross-level context **c**,

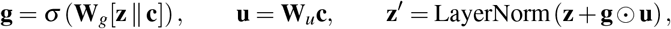

where [· ∥·] denotes concatenation, *σ* the sigmoid function, and ⊙ element-wise multiplication. The gate lets each node regulate how much cross-level information it incorporates. Residual connections between successive hierarchical layers stabilize optimization and preserve previously learned information. After repeated bidirectional propagation, every level encodes information from the full hierarchy while retaining its native structural resolution.

### 4.3 Model Training and Optimization

PRIME-ISP’s hierarchical GNN backbone and each task-specific output head use a hidden dimension of 128, 3 layers, and a dropout rate of 0.3; all tasks are trained with a batch size of 32. PRIME-ISP is trained end-to-end for each interaction class using a 5-fold cross-validation protocol. For a given task, the training set is partitioned into five folds with a fixed random seed; each fold trains an independent model on the remaining four folds, and predictions on the held-out test set are obtained by averaging the softmax outputs of all five fold models at inference time. Each fold is optimized with AdamW (learning rate 1 ×10^−3^, weight decay 1 ×10^−4^), preceded by a three-epoch linear warm-up from 1% to 100% of the target learning rate. After warm-up, the learning rate is reduced by a factor of 0.6 whenever the validation loss plateaus for five consecutive epochs (`ReduceLROnPlateau`). Gradients are clipped to a maximum norm of 5.0. Training runs for up to 200 epochs, with early stopping after 15 epochs (patience) without improvement in validation AUPRC.

Because interaction sites constitute a small minority of residues in every task, PRIME-ISP is trained with a combined objective that sums a class-weighted cross-entropy term and a soft Dice term,

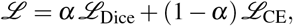

with *α* = 0.5. The cross-entropy term ℒ_CE_ is weighted by the inverse class frequency computed on the training set, penalizing misclassification of the minority interaction-site class more heavily. The Dice term

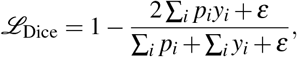

where *p*_*i*_ is the predicted probability of residue *i* belonging to the interaction-site class, *y*_*i*_ its ground-truth label, and *ε* a small smoothing constant, directly optimizes the overlap between predicted and true interaction sites and is largely insensitive to the class imbalance that dominates a per-residue cross-entropy loss. Combining the two terms lets the cross-entropy component maintain well-calibrated per-residue probabilities while the Dice component drives the model toward accurate site-level overlap. Model selection and early stopping are both driven by the area under the precision-recall curve (AUPRC) on a held-out validation fold.

### 4.4 Input and Output Adaptation for SHREC 2022 Evaluation

SHREC 2022 is designed as a purely geometric benchmark: to isolate pocket detection from any biological or chemical prior, the organizers anonymize each structure’s PQR file by substituting every atom name with the generic label “C”, every residue name with “UNK”, and randomly shuffling the order of all atom lines, discarding element identity, residue identity, and atom ordering alike^31^. PRIME-ISP’s hierarchical graph construction, in contrast, requires this biological information: element type and residue identity are used directly as handcrafted atomic and residue features (Section 4.2), and a consistent atom ordering is needed to recover backbone connectivity and assign atoms to their parent residues. To apply PRIME-ISP to this benchmark, we therefore align each anonymized PQR file back to its original, non-anonymized structure by matching atom coordinates, recovering atom names, residue identities, and a consistent atom order. This recovered information is used only to construct PRIME-ISP’s hierarchical graph and input features; the benchmark’s ground-truth binding-site labels, candidate-ranking protocol, and evaluation metrics are left unmodified. We note that this deviates from the strict geometry-only convention of the SHREC 2022 track, and our results are not intended as a direct, rule-consistent comparison against the track’s geometric baselines. Rather, our goal is to evaluate how much biological information, which PRIME-ISP is designed to exploit at every structural level, contributes to pocket identification on a benchmark where such information is deliberately withheld from the standard input. We note that this is not without precedent within the track itself: Fpocket, included by the organizers as an additional point of comparison, similarly uses full chemical information from the original PDB structures rather than the anonymized geometric representation.

The SHREC 2022 evaluation protocol scores predictions as spatially coherent, ranked binding pockets rather than independent per-atom or per-vertex classifications. We therefore add a post-processing step, specific to this benchmark, that converts PRIME-ISP’s atom-level predictions into a ranked set of candidate pockets. Given the atom-level interaction probabilities 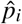 from the prediction head, we first convert each probability into a ligandability score,

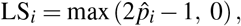

which rescales 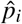 so that atoms with little evidence of being interacting contribute zero signal, rather than the small but nonzero weight a raw probability near 0.5 would otherwise carry. We discard atoms with 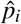 below a confidence threshold *T* = 0.5 and apply DBSCAN^39^ to the remaining atoms’ three-dimensional coordinates, grouping spatially proximal high-confidence atoms into candidate pockets and leaving isolated, low-confidence atoms unclustered. Each resulting cluster *C*_*k*_ is scored with a binding-region score,

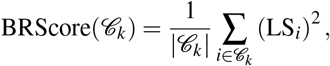

where squaring the ligandability scores emphasizes high-confidence predictions within a cluster, and averaging over cluster size so larger clusters aren’t favored purely for their extent. Candidate pockets are ranked by decreasing BRScore, and the ten highest-ranked clusters are retained, matching the benchmark’s evaluation protocol.

Surface-level evaluation requires an additional projection step because the molecular-surface representation used internally by PRIME-ISP does not preserve a one-to-one correspondence with the original SHREC 2022 mesh vertices. This reflects a limitation of the current framework. Although PRIME-ISP includes a molecular-surface level within its hierarchy and can, in principle, support surface-level prediction, the benchmark meshes contain substantially more vertices than the simplified surface representation used internally for efficient multiscale learning. Retaining the full-resolution benchmark mesh throughout hierarchical message passing would significantly increase the number of nodes and edges and, consequently, the computational and memory requirements of training and inference. We therefore predict interaction probabilities at the atomic level and subsequently project them onto the benchmark surface. To transfer these predictions, we build a KD-tree over the coordinates of the original SHREC mesh and, for each vertex, query its eight nearest protein atoms, weighting each atom’s contribution by the inverse of its distance to the vertex. The interaction probability of a surface vertex is then computed as the weighted average of these neighboring atom probabilities. Vertices whose nearest retained atom lies beyond 6 Å are considered uncovered by any prediction and assigned zero probability. This distance-weighted interpolation propagates atom-level predictions smoothly across the mesh while avoiding the artifacts that can arise from nearest-neighbor label assignment.

Because ligand-binding pockets are often associated with geometrically concave surface regions, we further incorporate local surface geometry into the pocket-ranking stage. Specifically, we compute the discrete mean curvature at each mesh vertex and define a concavity score by clipping the negative curvature to the interval [0, 1]. After clustering high-probability vertices with DBSCAN, we reweight the BRScore of each candidate pocket by a factor of 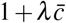, where 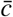 denotes the average concavity of the cluster, and *λ* = 0.5 controls the strength of the curvature-based boost. Surface-level candidate pockets are subsequently ranked using this curvature-adjusted score, and the top ten pockets are retained following the same protocol used for atom-level evaluation.

### 4.5 Evaluation Metrics

#### Classification metrics

Predictions are thresholded confidence scores compared against binary ground-truth labels, yielding true positives (TP), true negatives (TN), false positives (FP), and false negatives (FN). From these counts, we compute precision, recall, F1 score, and the Matthews correlation coefficient (MCC):

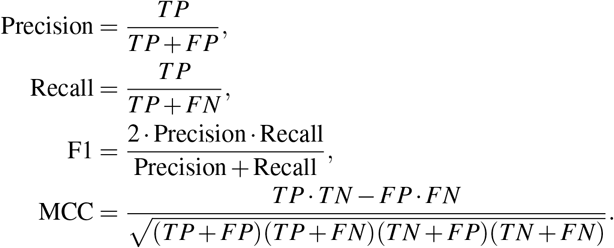

On the CarbPI benchmark, the same quantity is reported in the source literature as the Dice similarity coefficient (DSC), which is mathematically identical to F1; we report it as F1 throughout for consistency with the other interaction classes. We additionally report the area under the receiver operating characteristic curve (AUROC), computed by sweeping the classification threshold over the full range of predicted confidence scores, giving a threshold-independent summary of ranking quality. Because binding residues, atoms, and surface vertices are a minority class in every benchmark considered here, we treat MCC as the primary metric for comparing methods, since it remains informative under severe class imbalance where accuracy and AUROC can be dominated by the much larger negative class.

#### Pocket-ranking metrics

For benchmarks where predictions take the form of ranked candidate pockets rather than individual classifications, a candidate pocket is considered a hit for a given ligand if it satisfies both a ligand coverage (LC) and a pocket coverage (PC) threshold:

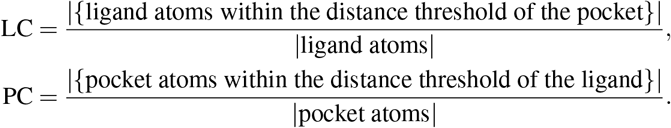

Following the benchmark’s own convention, the distance threshold is 5 Å when pockets are expressed as protein atoms and 4 Å when expressed as surface vertices. A hit is counted only when LC and PC both meet their respective thresholds (0.5 and 0.2). For each ligand, we determine its rank position as the number of higher-ranked candidate pockets that do not hit any ligand in the structure, so that pockets already matched to a different co-crystallized ligand do not penalize the ranking; a structure’s Top-*k* score is then the fraction of its ligands whose matching pocket falls within this rank position of *k*. Top1, Top3, and Top10 report this fraction averaged over all structures, for *k* = 1, 3, and 10 respectively, and the LC and PC values we report elsewhere are averaged over all successful hits.

## Data Availability

All datasets used in this study are publicly available at https://github.com/HySonLab/PRIME-ISP. This includes the benchmark partitions, ground-truth interaction-site annotations, and the case-study structures used in this work. Predicted protein structures generated with ESMFold 2 for benchmarks lacking an experimental structure are available on Zenodo at https://doi.org/10.5281/zenodo.23198572.

## Code Availability

The code used to train and evaluate PRIME-ISP, including data processing, model implementation, and analysis scripts, is publicly available at https://github.com/HySonLab/PRIME-ISP. Trained model checkpoints for all interaction classes are available on Zenodo at https://doi.org/10.5281/zenodo.23198572.

## Author Contributions

T.S.H. conceived the research idea. V.T.D.N. and T.S.H developed the PRIME-ISP framework and methodology. V.T.D.N implemented the experiments, and performed data analysis. V.T.D.N. wrote the initial manuscript draft. T.S.H. and J.K.J. supervised the project and provided critical feedback and revisions. All authors discussed the results and approved the final manuscript.

## Acknowledgments

This work used the Anvil system at Purdue University through allocation CIS260853, “GPU-Accelerated Foundation Models for Protein, Molecule, and Biomedical AI”, from the Advanced Cyberinfrastructure Coordination Ecosystem: Services & Support (ACCESS) program, which is supported by National Science Foundation grants #2138259, #2138286, #2138307, #2137603, and #2138296.

## References

1. Zhou, Y. et al. Ttd: Therapeutic target database describing target druggability information. Nucleic acids research 52, D1465–D1477 (2024).

2. Wakefield, A. E., Kozakov, D. & Vajda, S. Mapping the binding sites of challenging drug targets. Curr. opinion structural biology 75, 102396 (2022).

3. Qian, L. et al. The dawn of a new era: targeting the “undruggables” with antibody-based therapeutics. Chem. reviews 123, 7782–7853 (2023).

4. Shao, B., Li, P. & Liu, Z.-P. A protein surface-aware multimodal framework for residue-level metal-binding site recognition. Cell Reports Methods 6 (2026).

5. Zeng, M. et al. Protein–protein interaction site prediction through combining local and global features with deep neural networks. Bioinformatics 36, 1114–1120 (2020).

6. Li, Y., Golding, G. B. & Ilie, L. Delphi: accurate deep ensemble model for protein interaction sites prediction. Bioinformatics 37, 896–904 (2021).

7. Mou, M. et al. A transformer-based ensemble framework for the prediction of protein–protein interaction sites. Research 6, 0240 (2023).

8. Fang, Y. et al. Deepprosite: structure-aware protein binding site prediction using esmfold and pretrained language model. Bioinformatics 39, btad718 (2023).

9. Petsalaki, E., Stark, A., García-Urdiales, E. & Russell, R. B. Accurate prediction of peptide binding sites on protein surfaces. PLoS computational biology 5, e1000335 (2009).

10. Zhao, Z., Peng, Z. & Yang, J. Improving sequence-based prediction of protein–peptide binding residues by introducing intrinsic disorder and a consensus method. J. Chem. Inf. Model. 58, 1459–1468 (2018).

11. Wang, R., Jin, J., Zou, Q., Nakai, K. & Wei, L. Predicting protein–peptide binding residues via interpretable deep learning. Bioinformatics 38, 3351–3360 (2022).

12. Zhu, Y.-H., Hu, J., Song, X.-N. & Yu, D.-J. Dnapred: accurate identification of dna-binding sites from protein sequence by ensembled hyperplane-distance-based support vector machines. J. chemical information modeling (2019).

13. Liu, R. & Hu, J. Dnabind: a hybrid algorithm for structure-based prediction of dna-binding residues by combining machine learning-and template-based approaches. PROTEINS: structure, Funct. Bioinforma. 81, 1885–1899 (2013).

14. Lam, J. H. et al. A deep learning framework to predict binding preference of rna constituents on protein surface. Nat. communications 10, 4941 (2019).

15. Xia, Y., Xia, C.-Q., Pan, X. & Shen, H.-B. Graphbind: protein structural context embedded rules learned by hierarchical graph neural networks for recognizing nucleic-acid-binding residues. Nucleic acids research 49, e51–e51 (2021).

16. Zhang, J. et al. Deeppanb: integrating protein language model with painn equivariant graph neural networks for prediction of protein–nucleic acid binding sites. Briefings Bioinforma. 27, bbag450 (2026).

17. Canner, S. W., Shanker, S. & Gray, J. J. Structure-based neural network protein–carbohydrate interaction predictions at the residue level. Front. Bioinforma. 3, 1186531 (2023).

18. Le Guilloux, V., Schmidtke, P. & Tuffery, P. Fpocket: an open source platform for ligand pocket detection. BMC bioinformatics 10, 168 (2009).

19. Krivák, R. & Hoksza, D. P2rank: machine learning based tool for rapid and accurate prediction of ligand binding sites from protein structure. J. cheminformatics 10, 39 (2018).

20. Mylonas, S. K., Axenopoulos, A. & Daras, P. Deepsurf: a surface-based deep learning approach for the prediction of ligand binding sites on proteins. Bioinformatics 37, 1681–1690 (2021).

21. Pham, P., Nguyen, V. T. D., Song, K., Chen, J. & Hy, T.-S. Linker: L earning in teractions between functional groups and residues with chemical k nowledge-e nhanced r easoning and explainability. J. Chem. Inf. Model. 66, 8579–8593 (2026).

22. Wu, Q. et al. Decoupling topological and molecular features for interpretable biomolecular interaction prediction. Briefings Bioinforma. 27, bbag471 (2026).

23. Mou, M. et al. Accurate identification of protein binding sites for all drug modalities using allsites. Adv. Sci. 13, e16530 (2026).

24. Nguyen, V. T. D., Johnstone, J. K. & Hy, T.-S. Prime: Protein representation via physics-informed multiscale equivariant hierarchies. In International Workshop on Data Mining in Bioinformatics at ACM SIGKDD (2026). Accepted.

25. Gainza, P. et al. Deciphering interaction fingerprints from protein molecular surfaces using geometric deep learning. Nat. methods 17, 184–192 (2020).

26. Zhao, A. et al. Surf2spot: a surface-informed geometry-aware model for predicting partner-independent binder and nanobody design hotspots. Briefings Bioinforma. 27, bbag542 (2026).

27. Candido, S. et al. Language modeling materializes a world model of protein biology (2026). Preprint.

28. Wang, S., Chen, W., Han, P., Li, X. & Song, T. Rgn: residue-based graph attention and convolutional network for protein–protein interaction site prediction. J. chemical information modeling 62, 5961–5974 (2022).

29. Mahbub, S. & Bayzid, M. S. Egret: edge aggregated graph attention networks and transfer learning improve protein–protein interaction site prediction. Briefings Bioinforma. 23, bbab578 (2022).

30. Zheng, M., Sun, G., Li, X. & Fan, Y. Egpdi: identifying protein–dna binding sites based on multi-view graph embedding fusion. Briefings Bioinforma. 25, bbae330 (2024).

31. Gagliardi, L. et al. Shrec 2022: protein–ligand binding site recognition. Comput. & Graph. 107, 20–31 (2022).

32. Decherchi, S. & Rocchia, W. A general and robust ray-casting-based algorithm for triangulating surfaces at the nanoscale. PloS one 8, e59744 (2013).

33. Zhao, H., Jiang, L., Jia, J., Torr, P. H. & Koltun, V. Point transformer. In Proceedings of the IEEE/CVF international conference on computer vision, 16259–16268 (2021).

34. Yu, H. et al. Aptameric hirudins as selective and reversible exosite-active site (exact) inhibitors. Nat. Commun. 15, 3977 (2024).

35. Meador, W. E., Means, A. R. & Quiocho, F. A. Target enzyme recognition by calmodulin: 2.4 å structure of a calmodulin-peptide complex. Science 257, 1251–1255 (1992).

36. Schrödinger, LLC. The PyMOL molecular graphics system, version 1.8 (2015).

37. Kabsch, W. & Sander, C. Dictionary of protein secondary structure: pattern recognition of hydrogen-bonded and geometrical features. Biopolym. Orig. Res. on Biomol. 22, 2577–2637 (1983).

38. Lin, Z. et al. Evolutionary-scale prediction of atomic-level protein structure with a language model. Science 379, 1123–1130 (2023).

39. Ester, M., Kriegel, H.-P., Sander, J., Xu, X. et al. A density-based algorithm for discovering clusters in large spatial databases with noise. In kdd, vol. 96, 226–231 (1996).

